# Unveiling the Diversity and Demography of Sharks and Rays at key landing sites in Bangladesh

**DOI:** 10.1101/2025.06.26.660850

**Authors:** Fahmida Khalique Nitu, Dilshad Farahnaz Tasnim, Sultan Ahmed, Md Kutub Uddin, Shaili Johri

**Author notes:** Corresponding Author Information: Fahmida Khalique Nitu Shaili Johri.

## Abstract

In Bangladesh’s marine waters in the Bay of Bengal, sharks and rays face grave threats from unsustainable fishing practices and habitat degradation. Findings from our taxonomic assessment of elasmobranchs landed at the two most important marine fish landing stations in Bangladesh bring new information and insights about the nature of the Threatened elasmobranch bycatch. This study presents findings from a taxonomic assessment at Cox’s Bazar and Chattogram, two crucial landing stations in Bangladesh where most shark bycatch landings happen. Over two years, 525 fish were sampled; 206 samples were of sharks, 91 of rays, and 64 of rhino rays, representing over 29 species across 12 families. Among them are 4 families of sharks, 5 rays, and 3 rhino rays. The predominant families observed were *Carcharhinidae, Glaucostegidae*, and *Rhinopteridae*. The most frequently landed species were *Sphyrna lewini, Glaucostegus younholeei, Carcharhinus leucas, Gymnura poecilura, and Mobula mobular.* Out of the 525 samples, 65% (n=340) were female, 35% (n=183) were male, and the rest (n=2) were unidentified. Finding one sample of *Pristis pristis* indicates the severity of the decline. Additionally, for *Glaucostegus granulatus,* 12 among the 13 studied specimens were adult females. A significant proportion of the sampled specimens, 45%, were juveniles, while only 24% were adults. This is likely because these fishing boats bringing the catch operate in inshore water and nearshore seas, which are potential nursery grounds. The overlapping of fishing grounds and these critical habitats poses a significant challenge in elasmobranch conservation and hinders population recovery. The high number of juveniles and female adults in the catch reminds us of the necessity of integrating threatened bycatch reduction measures into the country’s fisheries management.

## 1. Introduction

Sharks and rays (hereafter referred to as elasmobranchs) play a crucial role in maintaining ocean health by regulating population dynamics across various trophic levels (Ferretti et al., 2008; Stevens et al., 2000). As apex predators in the marine food web, they help control ocean ecosystems through top-down regulation (Dedman et al., 2024; Frisk et al., 2005). Some species of elasmobranchs, such as the narrow sawfish and large-tooth sawfish, also inhabit freshwater and estuarine ecosystems (Jabado et al., 2017). The ecological significance of elasmobranchs underscores the urgent need for their conservation. Their slow growth rate, low fecundity, and lengthy gestation period make them particularly vulnerable to fishing mortality (Cortés, 2000). Elasmobranchs face considerable threats, including overexploitation from mostly bycatch and in some cases targeted fishing, which have led to significant population declines. Extensive bycatch and targeted fishing of rays are major concerns, as they contribute to the decrease in elasmobranch populations (Dulvy et al., 2014). A significant portion of elasmobranch bycatch is discarded at sea and remains unreported (Davidson et al., 2016). While some fishers retain valuable bycatch, the amount of retained catch varies depending on the species and their economic value to the fishers.

Fishing pressure on elasmobranchs is increasing globally, driven by the rising demand for meat, fins, and liver oil (Fowler et al., 2005). Fins are highly prized, especially in certain Asian countries where shark fin soup is considered a delicacy (Clarke et al., 2006). Artisanal fisheries globally also report substantial juvenile landings (Appleyard et al., 2018). In addition to overfishing, coastal habitat degradation, pollution, infrastructure development, and climate change are significant factors contributing to the decline of these species (Jabado et al., 2017).

Among elasmobranchs, species from the sawfish, wedgefish, and guitarfish families are globally threatened (Davidson et al., 2016; Jabado et al., 2017). Once common in the Indian Ocean region, sawfish are now extremely rare, with recent reports indicating only a few occurrences in the northeastern Arabian Sea (Jabado et al., 2017). These species are highly vulnerable to gillnets and other demersal trawl nets, highlighting the need for targeted conservation efforts in this region.

Elasmobranchs are the first major marine fish lineage at risk of extinction on a global scale (Dulvy et al., 2021). According to the most recent IUCN assessment, 37% of elasmobranchs are categorized as critically endangered, endangered, or vulnerable. The situation is even more dire in the Arabian Ocean region (Jabado et al., 2017). Traditional management measures for other fish species may not be effective for elasmobranchs due to their distinct life history traits. Therefore, a specialized management system for elasmobranchs is essential at local, national, and regional levels.

Elasmobranch fisheries are vital to Asia’s marine ecosystems and economic landscape. The region is home to some of the largest populations of sharks and rays globally, with many species targeted for their fins, meat, liver oil, and other products. The Indian Ocean is considered a hotspot for elasmobranch diversity (Jaquemet et al., 2023; Roberts et al., 2002). Southeast Asia and the Bay of Bengal play a significant role in global shark and ray fisheries, supporting both artisanal and international commercial markets (Haque et al., 2021). These areas’ diverse habitats and high productivity support dense coastal communities that are integral to the global seafood supply chain. However, overfishing has led to substantial elasmobranch bycatch, underscoring the region’s importance in elasmobranch exports. Effective fisheries management in this area is crucial for preserving biodiversity, supporting local livelihoods, and ensuring the sustainable use of marine resources.

Fishing mortality of sharks and rays in Bangladesh is considerable, and the country contributes significantly to regional shark fisheries in Asia (Haque et al., 2021). With its extensive coastline and rich marine biodiversity, Bangladesh is known for its significant elasmobranch bycatch and trade (DoF, 2020). The fisheries sector in Bangladesh supports both domestic food security and the global seafood trade (Hoq et al., 2011). However, sharks and rays face severe threats from unsustainable fishing practices, habitat degradation, and inadequate management, leading to alarming population declines (Stevens et al., 2000). As a result, many species in the Bay of Bengal are now classified as threatened or endangered by the IUCN (Fowler et al., 2013).

Elasmobranch fishing grounds in Bangladesh are primarily located in the southwestern region, offshore of Kuakata, Dublar Char, Patharghata, and Barguna, as well as in the southeastern region off the shores of Kutubdia, Moheskhali, Cox’s Bazar, Teknaf, and Swandip (Hoq et al., 2011). Primary landing sites for elasmobranchs in Bangladesh include Cox’s Bazar, Chattogram, Mohipur, and Patharghata, where commercial fishing boats bring their catches to the landing stations. Despite their low contribution to the mainstream fisheries sector, elasmobranch species in Bangladesh have limited research, poor documentation, and inadequate data availability (Badhon et al., 2019; Haque et al., 2021).

In Bangladesh, elasmobranch catches predominantly result from bycatch alongside other commercially targeted fish species (Badhon et al., 2019; Hoq et al., 2011). There are 220 large-scale industrial trawlers and 67,669 commercial boats operating with various fishing gears (Badhon et al., 2019; DoF, 2020). The fishing vessels use multiple gears such as gillnets, set bag nets, bottom longlines, beach seines, and trammel nets, targeting species like *Eleutheronema tetradactylum* (Fourfinger Threadfin), *Tenualosa Ilisha* (Hilsha Shad), *Auxis rochei* (Bullet Tuna), *Harpadon nehereus* (Bombay Duck), and *Pampus argenteus* (Silver Pomfret). Only a few small-scale commercial boats specifically target rays, often using hooks (Roy et al., 2015). According to fisheries statistics, significant bycatch is caught primarily through gillnet and longline fishing (DoF, 2020). However, many elasmobranchs are discarded at sea and never reported. Fishers tend to prioritize keeping batoids (rays) from bycatch to sell, due to the demand for products like skins, which provide additional income.

Bangladesh is a signatory to the Convention on International Trade in Endangered Species of Wild Fauna and Flora (CITES) and the Convention on Biological Diversity (CBD), both of which require commitments toward sustainable biodiversity management, including the conservation of elasmobranch species. While Bangladesh’s fisheries laws do not include bycatch reduction measures for elasmobranchs, the country has enacted “The Bangladesh Wildlife (Conservation and Security) Act 2012,” which provides a legal framework for CITES implementation. This act prohibits the catching and trading of 29 elasmobranch species under two schedules. However, enforcement is challenging, as elasmobranchs often come as bycatch in otherwise legal fishing operations.

Overall, studying elasmobranchs will support marine conservation efforts in Bangladesh by identifying key threats and enabling better management strategies. It will also contribute to sustainable fisheries that support local livelihoods and the regional economy, while helping Bangladesh meet its international commitments and national laws to protect endangered species. As an important hub for elasmobranchs in Southeast Asia, understanding the dynamics of elasmobranch fisheries is critical for ocean health and the sustainability of coastal livelihoods.

This study presents findings on seasonal diversity at two major fishing sites in Bangladesh: Chattogram and Cox’s Bazar. These sites are the leading contributors to elasmobranch landings among the 19 coastal countries in the region. We identify and describe the morphology of 29 landed species. Bangladesh is one of the most data-deficient countries regarding elasmobranch species (Haque et al., 2021). This study aims to improve our understanding of elasmobranch diversity and landing morphology in the Bay of Bengal. It seeks to update taxonomic information, fill existing knowledge gaps at key sites, and provide data that will inform effective conservation measures. Additionally, documenting elasmobranch bycatch will offer valuable insights into the diversity of these species in Bangladesh and the impact of fisheries on their populations.

## 2. Methods

### 2.1. Data Collection

The current study was conducted in the two largest and high-volume landing stations in the southeastern part of Bangladesh (**Figure 1**). The sites are the Bangladesh Fisheries Development Cooperation (BFDC) operated landing stations in Cox’s Bazar and Chattogram. Cox’s Bazar landing station has a designated location for elasmobranch landings, while in Chattogram, they were collected by three or four specific shops after being purchased from fishers by the traders.

**Figure 1:**
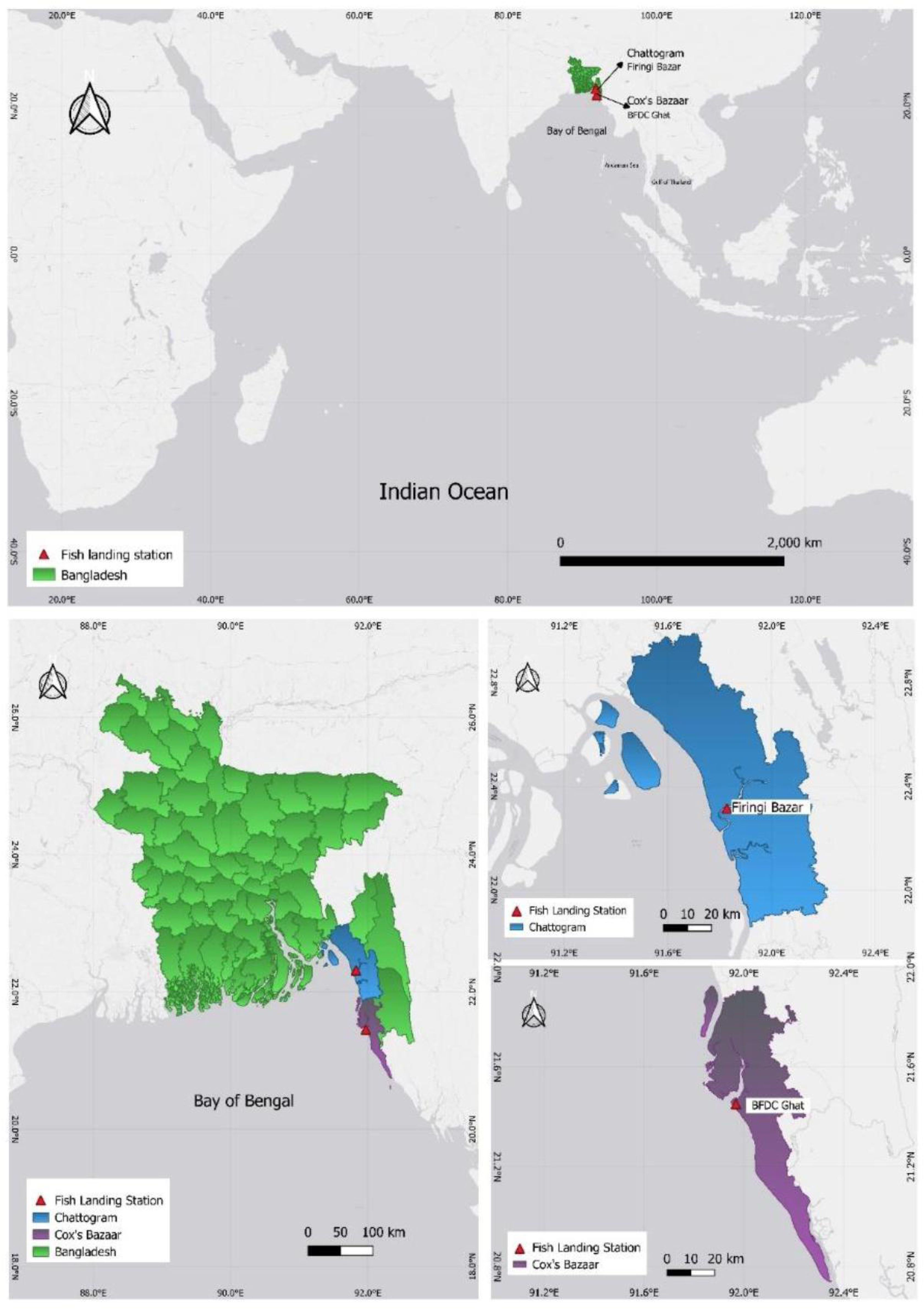
Sampling site located along the coast of Bay of Bengal under the Greater Indian Ocean. The two-sampling site – Chattogram and Cox’s Bazar were selected due to high volume of Elasmobranch landings in this region.

These two sites have a higher volume of landings compared to other coastal sites in Bangladesh. They are the home ports of most commercial boats and large-scale industrial trawlers, or the business operations of the owners based here. Also, these two sites have safe harbors for many vessels. Moreover, most traders and processing plants for shark products are located around these areas. Two categoric data (length and width) and two numeric data (sex and life stage) were collected during the period.

### 2.2. Study duration

During the initial phase of our study, we collected data from Cox’s Bazar between February and April 2022 and from Chattogram between March and May 2022. We chose to conduct our study during the winter and summer months because these periods experience lower rainfall and more favorable weather conditions, allowing fishermen to engage in deep-sea fishing activities. We conducted sampling during the full moon and new moon periods of each month. Due to the gravitational forces, spring tides occur, and nutrients stir up during these periods, increasing fish assemblage in certain areas. These are typically times when fish landings are high.

### 2.3. Morphological identification

Photographs of both the dorsal and ventral sides of the fish, as well as close-up shots of the fins, were taken. However, in some cases, traders were unwilling to flip the fish, so photographs were taken of only one side. Along with that, photographic identification was also done following the standard protocols (Ebert, Ho, White, & De Carvalho, 2013; Last et al., 2016). Species identifications were verified by utilizing the citizen science species identification platform iNaturalist.org. All the photographs were uploaded to the database ‘iNaturalist’, and expert observers’ identification was documented on the datasheet along with our taxonomic identification. Informal interviews with fishers and traders helped to understand the gear type, trip lengths, and elasmobranch bycatch percentage.

### 2.4. Maturity and Sex determination

We measured the samples at the landing sites both manually and digitally. All measurements were then converted to centimeters. Total length (TL) was measured for sharks and rhinorays, while ray’s disc width (DW) was measured. Average maturity length and width for males and females were determined using existing literature (**Table 1**).

**Table 1:**
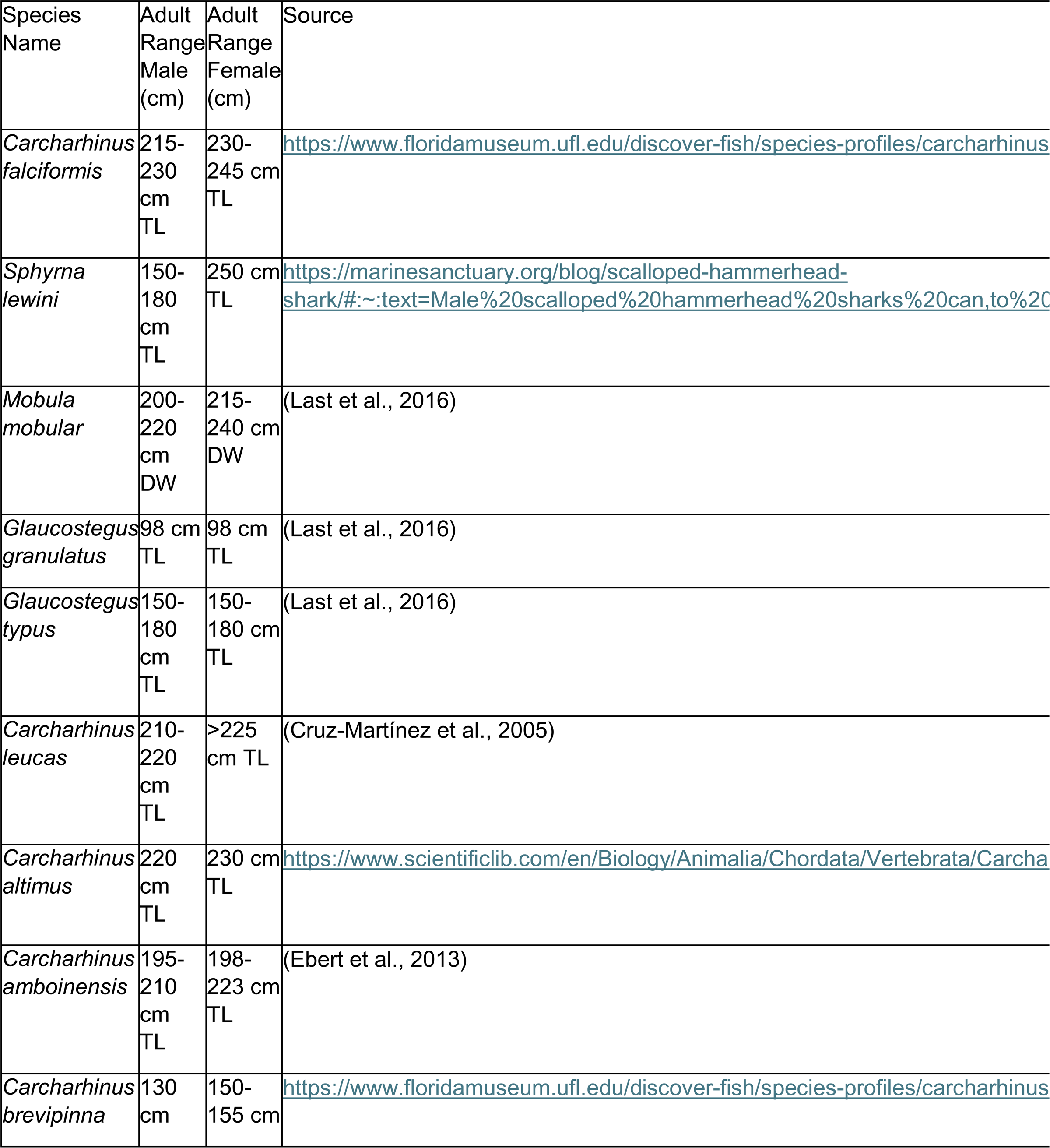
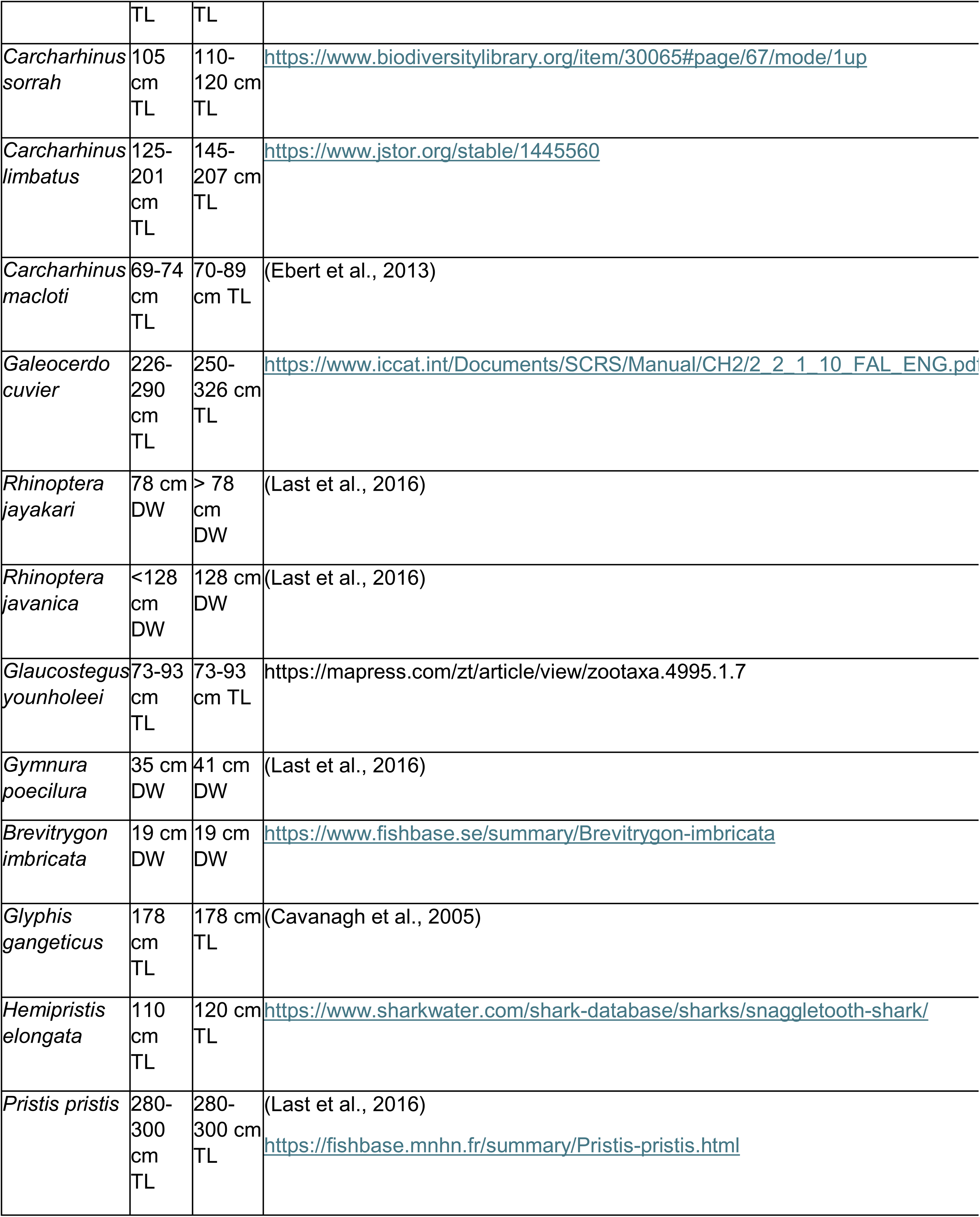
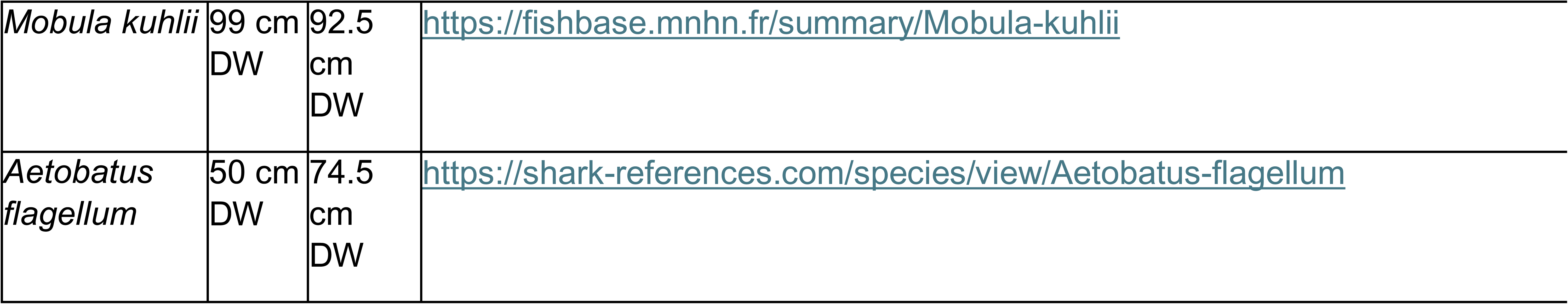
Total length (TL) and Disc Width (DW) (cm) threshold used to determine maturity for each species are listed below.

Sex was determined in the field by observing the presence of claspers. However, samples were identified at the lowest possible taxonomic level to determine the species composition of the landings (Ebert et al., 2013; Jabado & Ebert, 2015; Kottillil et al., 2023). Length and width measurements were later used to identify the life stage or maturity by following IUCN Red List guidelines and other relevant kinds of literature (Ebert et al., 2013).

### 2.5. Conservation status

The conservation status of species in Bangladesh was evaluated using the IUCN Red List of Threatened Species. We did this to determine the impact of fisheries on the priority species. The IUCN conservation categories were used, which include Critically Endangered (CE), Endangered (EN), Vulnerable (VU), Near Threatened (NT), Least Concern (LC), and Data Deficient (DD).

## 3. RESULTS

### 3.1. Species Distribution

We conducted a survey and sample collections from Bangladesh’s two biggest marine fish landing stations and opportunistically sampled 525 elasmobranchs (**Figure 2).** During the project’s first phase in 2022, we photographed, measured, and collected 83 specimens from Chattogram and 145 specimens from Cox’s Bazar. In the second phase of our study, in 2023, we conducted a survey involving the photographing and measuring of 117 specimens from Chattogram and 180 specimens from Cox’s Bazar. In total, 525 elasmobranch specimens were collected opportunistically, encompassing two superorders: **Selachimorpha** and **Batoidea**. The survey resulted in approximately 2,000 photographs of elasmobranchs.

**Figure 2:**
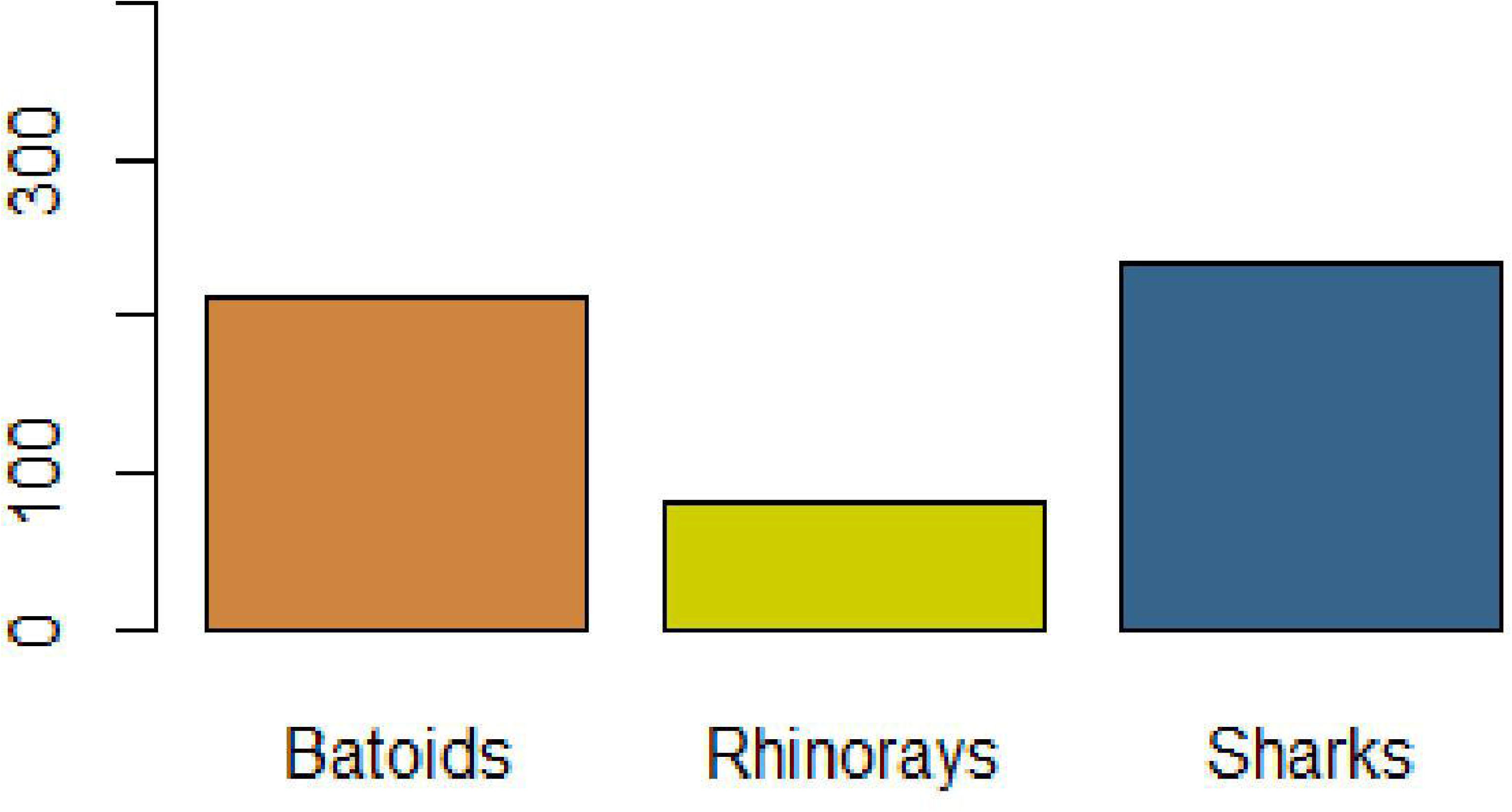
Number of Elasmobranch by three categories of Batoids Rhinorays and Sharks across the two landing stations Chattogram and Cox’s Bazar.

Our specimen collection included 12 species from the Superorder Selachimorpha, which comprised 1 order, 4 families, and 5 genera (**Figure 3A**, **Table 2)**. *Sphyrna lewini* (Scalloped Hammerhead) was the most frequently landed shark species in Cox’s Bazar and Chattogram, followed by *Carcharhinus leucas* (Bull shark) (**Figure 3A**, **Table 2)**. A total of 87 *Sphyrna lewini* were collected.

**Figure 3:**
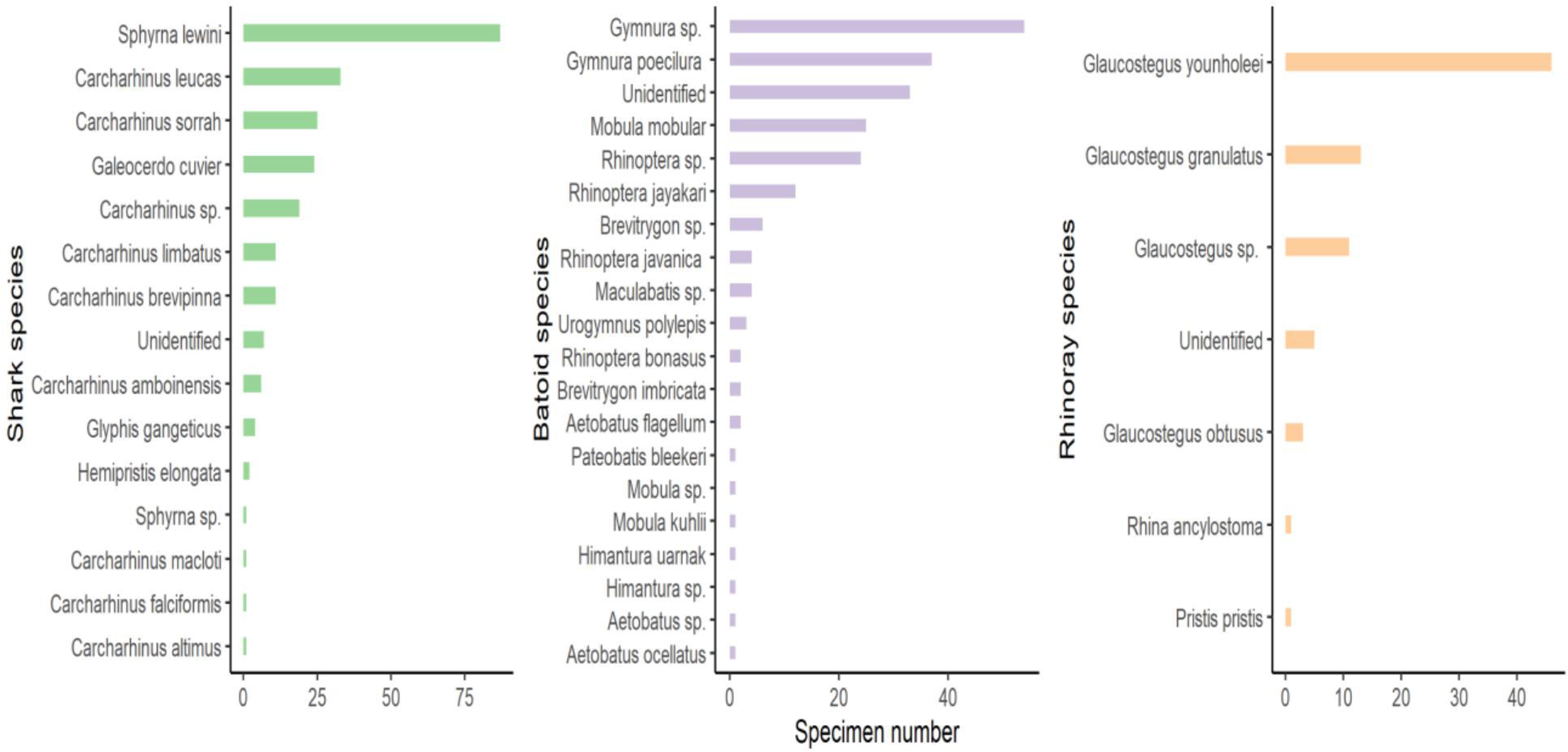
Specimen number of each Shark, Batoid and Rhinoray species.

Specimens in the Superorder Batoidea comprised 17 species organized into three orders, eight families, and 11 genera. *Gymnura poecilura* (Long tail Butterfly Ray) exhibited the highest abundance (n=37) among the ray species in number, followed by *Mobula mobular* (Spinetail Devil Ray) (**Figure 3B**, **Table 2)**. Among the Rhinorays, *Glaucostegus younholeei* (Bangladeshi Guitarfish) was the highest in number, with a total of 46 samples collected (**Figure 3C**, **Table 2)**.

In total, 12 species of sharks (**Figure 4)**, 12 species of rays (**Figure 5)**, and 5 species of rhino rays (**Figure 6)** were observed in the study. We avoided sampling the *Scoliodon* species group because it was one of the most abundant and frequently sighted species (Haque et al., 2022) and Near threatened species according to IUCN (Kyne et al., 2020). For each species, the images used for species identification and the process of research-grade identifications through iNaturalist can be found at the links provided in **Supplementary Table 3**.

**Figure 4:**
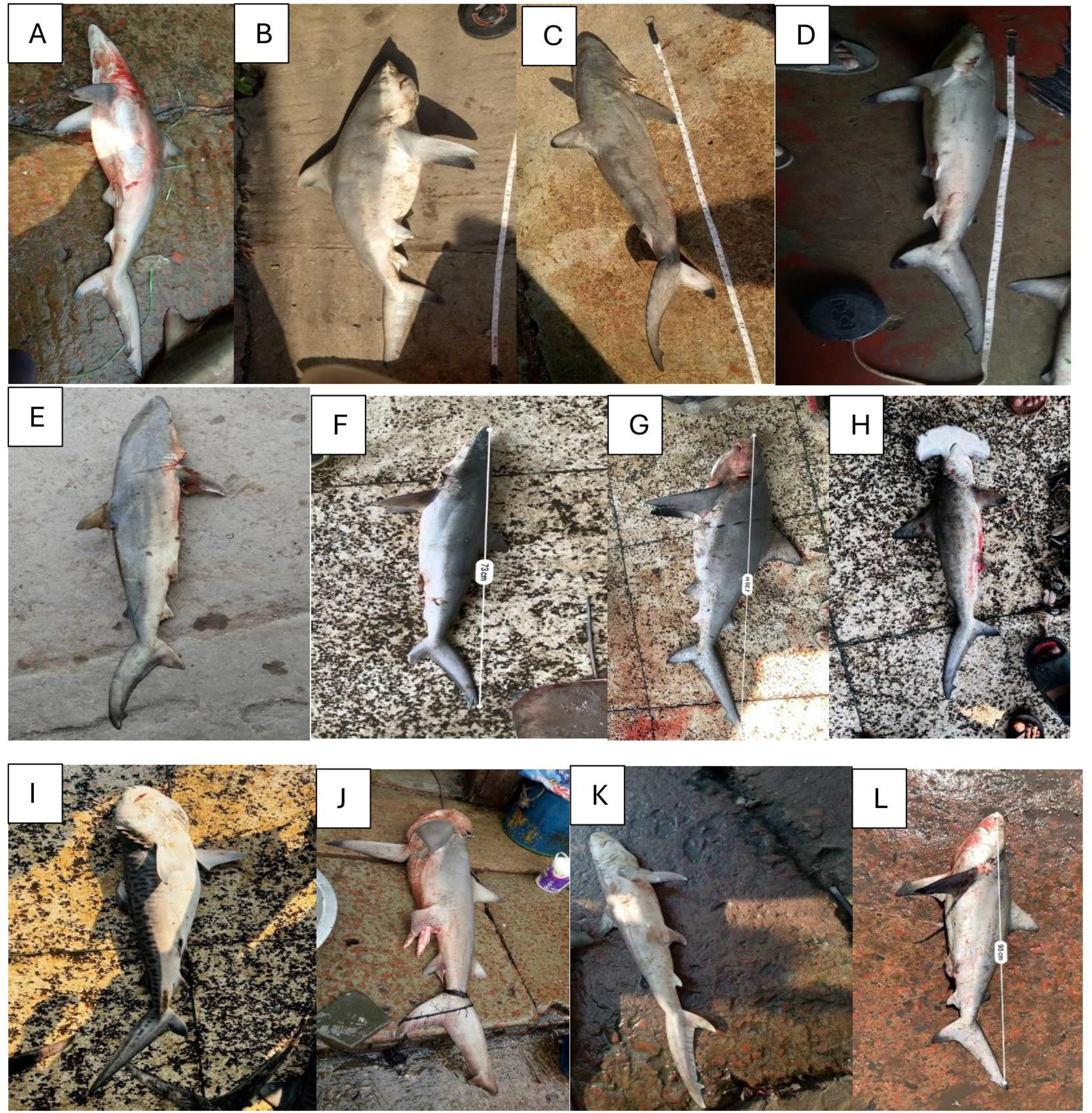
Representative pictures of species sampled in the superorder Selachimorpha. (**A)** *Carcharhinus brevipinna* (Spinner Shark), (**B)** *Carcharhinus leucas* (Bull Shark), (**C)** *Carcharhinus limbatus* (Blacktip Shark), (**D)** *Carcharhinussorrah* (Spot-tail Shark), (**E)** *Carcharhinus macloti* (Hardnose Shark) (**F)** *Carcharhinus falciformis* (Silky Shark), (**G)** *Carcharhinus amboinensis* (Pigeye Shark) (**H)** *Sphyrna lewini* (Scalloped Hammerhead Shark) (**I)** *Galeocerdo cuvier* (Tiger Shark) (**J)** *Glyphis gangeticus* (Ganges River Shark) (**K)** *Hemipristis elongata* (Snaggletooth Shark) (**L)** *Carcharhinus altimus* (Bignose Shark).

**Figure 5:**
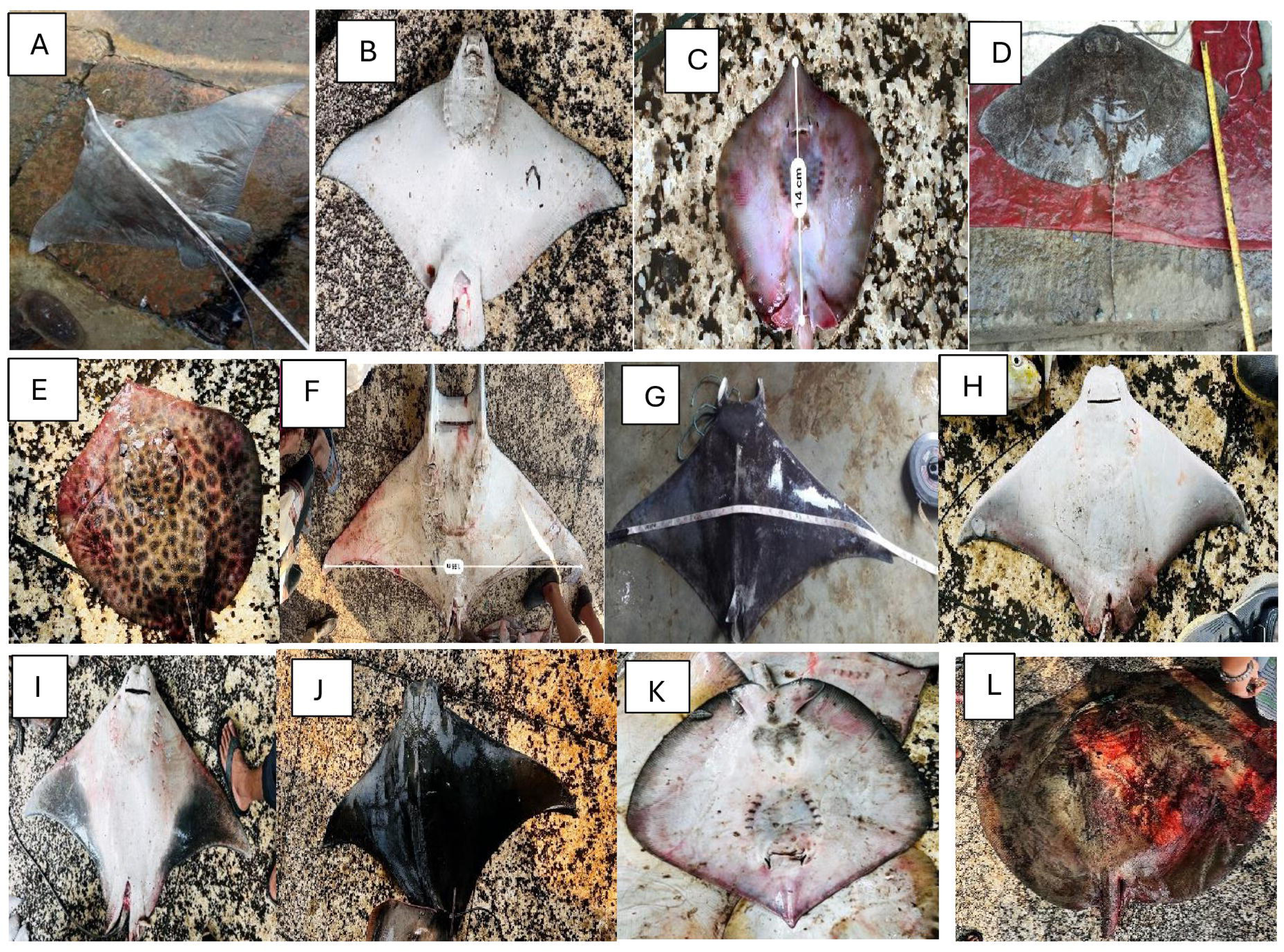
Representative pictures of species sampled in the superorder Batoidea. (**A)** *Aetobatus flagellum* (Longheaded Eagle Ray), (B) *Aetobatus ocellatus* (Ocellated Eagle Ray), (**C)** *Brevitrygon imbricata* (Bengal Whipray), (D) *Gymnura poecilura* (Longtail Butterfly Ray), (**E)** *Himantura uarnak* (Honeycomb Stingray) (**F)** *Mobula mobular* (Spine-tail Devil Ray), (**G)** *Mobula kuhlii* (Short-horned Pygmy Devil Ray) (**H)** *Rhinoptera bonasus* (Cownose Ray) (**I)** *Rhinoptera javanica,Javanese* Cownose Ray) (**J)** *Rhinoptera jayakari* (Short-tail Cownose Ray) (**K)** *Pateobatis bleekeri* (Bleeker’s Whipray) (**L)** *Urogymnus Polylepis* (Giant Freshwater Sting Ray).

**Figure 6:**
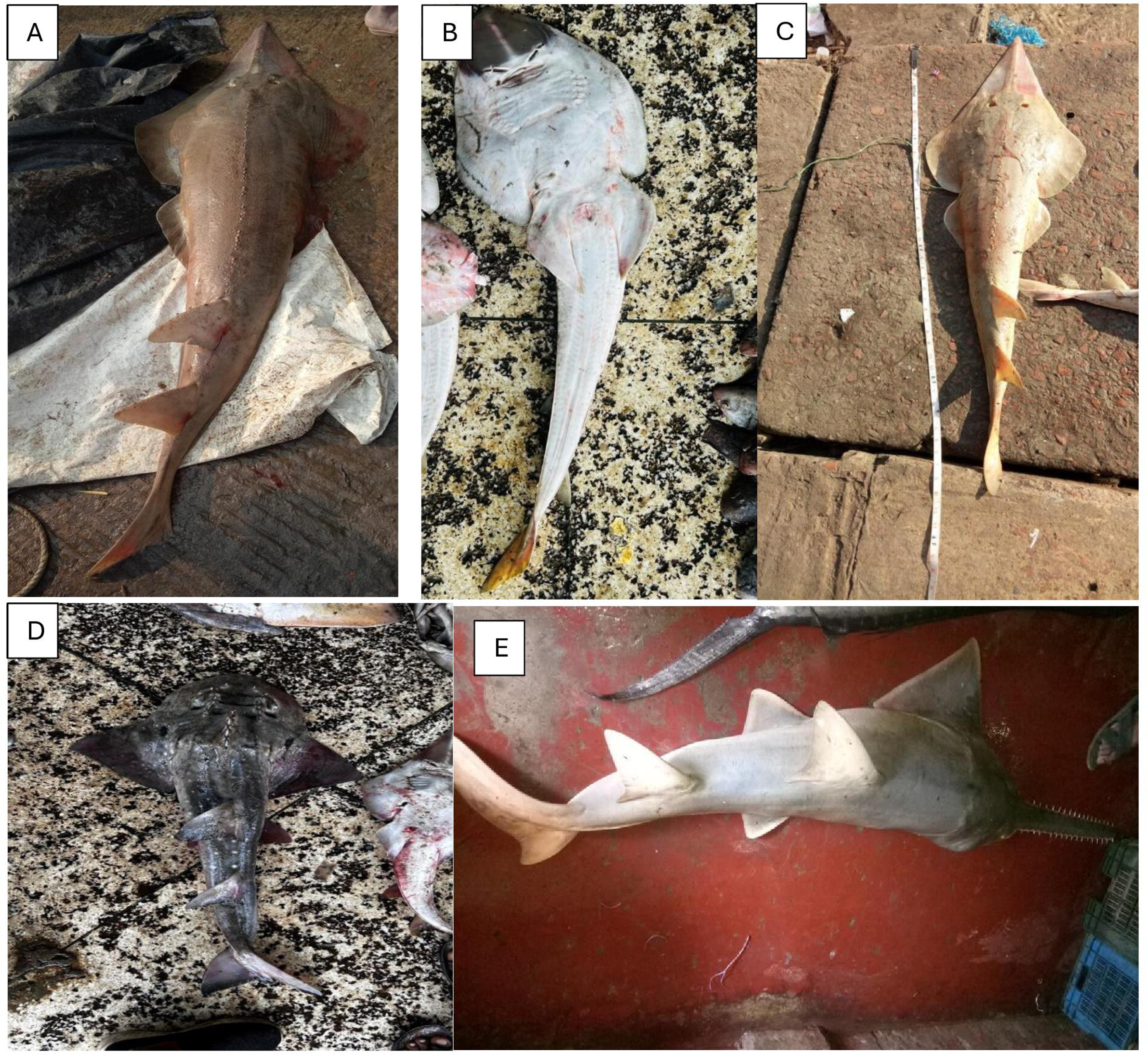
Representative pictures of species sampled in the superorder Batoidea. (**A)** *Glaucostegus granulatus* (Sharpnose Guitarfish), (B) *Glaucostegus obtusus* (Widenose Guitarfish), (**C)** *Glaucostegus younholeei* (Bangladeshi Guitarfish), (**D)** *Rhina ancylostoma* (Bowmouth Guitarfish), (E) *Pristis pristis* (Largetooth Sawfish).

We could not determine the taxonomic identification of 163 samples beyond the genus or family level, and these are not included in **Table 2** because photographs of some samples were not well captured, and some samples were dismembered with inadequate morphological information.

The two sampling sites differed in the species type and abundance of the species sampled. Cox’s Bazar samples were dominated by *Sphyrnidae* (20%), followed by *Carcharhinidae* (20%) and *Gymnuridae* (15%). Whereas Chattogram samples were dominated by *Carcharhinidae* (27%), *Gymnuridae* (24%), *Glaucostegidae* (19%), and *Sphyrnidae* (11%). Frequently landed families of rays were *Gymnuridae*, *Mobulidae*, *Mylobatidae*, *Glaucostegiade*, *Rhinopteriade*, *Dasyiatidae*.The least abundant species sampled were *Pristis pristis* (Largetooth Sawfish) and *Glyphis gangeticus* (Ganges Shark). Only one specimen of *Pristis pristis* and four specimens of *Glyphis gangeticus* were sampled in Chattogram. The sampling of different species at the two locations may indicate distinct distribution patterns for the respective species in the fishing grounds used by fishers landing their catch at the two sampling sites, and by extension, distinct ecology and diverse habitats in the Bay of Bengal region.

### 3.2. Conservation Status

Elasmobranch species were identified across all conservation categories in the two sampling locations. Of the 29 species of our study 24.13% are Critically Endangered, 27.58% are Endangered, 34.48% are Vulnerable, and 13.79% are Near Threatened.

The critically endangered species among our samples were *Sphyrna lewini* (n=87) (Scalloped Hammerhead), *Glyphis gangeticus* (n=4) (Ganges River Shark), *Pristis Pristis* (n=1) (Largetooth Sawfish), *Glaucostegus granulatus* (n=13) (Granulated Guitarfish), *Glaucostegus younholeei* (n=46) (Bangladeshi Guitarfish), *Glaucostegus obtusus* (n=3) (Widenose Guitarfish), *and Rhina ancylostomus* (n=1) (Bowmouth Guitarfish) (**Figure 7A)**. Endangered species among our samples include *Aetobatus flagellum* (Loggerheaded Eagle Ray), *Himantura uarnak* (Honeycomb Stingray), *Mobula mobular* (Spinetail Devil Ray), *Mobula kuhlii* (Shortfin Devil Ray), *Rhinoptera javanica* (Javanese Cownose ray), *Rhinoptera jayakari* (Oman Cownose ray), *Pateobatis bleekeri* (Bleekers Whipray), *Urogymnus Polylepis* (Giant Freshwater Stingray) and vulnerable species included *Carcharhinus brevipinna* (Spinner Shark), *Carcharhinus leucas* (Bull Shark), *Carcharhinus limbatus* (Common Blacktip Shark), *Carcharhinus falciformis* (Silky shark), *Carcharhinus amboinensis* (Pigeye Shark), *Hemipritis elongata* (Snaggletooth Shark), *Aetobatus ocellatus* ( Whitespotted Eagle Ray), *Brevitrygon imbricata* (Scaly Whip Ray), *Gymnura poecilura* (Longtail Butterfly Ray), *Rhinoptera bonasus* (Cownose Ray) (**Figure 7A)** and near threatened were *Carcharhinus sorrah* (Spottail Shark), *Carcharhinus macloti* (Hardnose Shark), Galeocerdo cuvier (Tiger shark), *Carcharhinus altimus* (Bignose Shark).

**Figure 7A:**
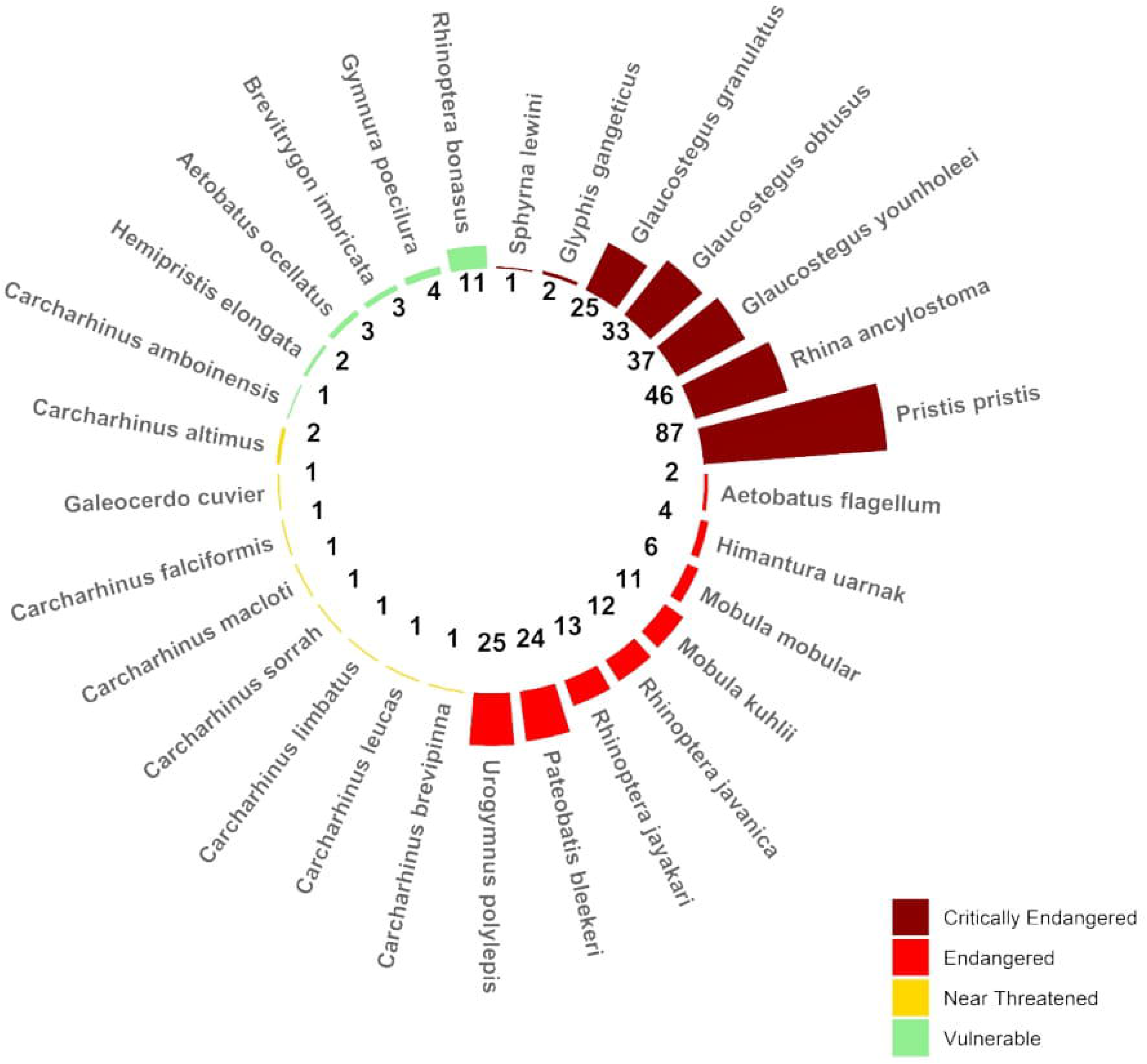
Species composition of sharks from Chattogram and Cox’s Bazar during the study period. Species are listed and grouped according to frequency. IUCN Red List status is given for each species and colour coded. Here Critically Endangered (dark red), Endangered (red), Near Threatened (yellow), Vulnerable (green).

**Figure 7B:**
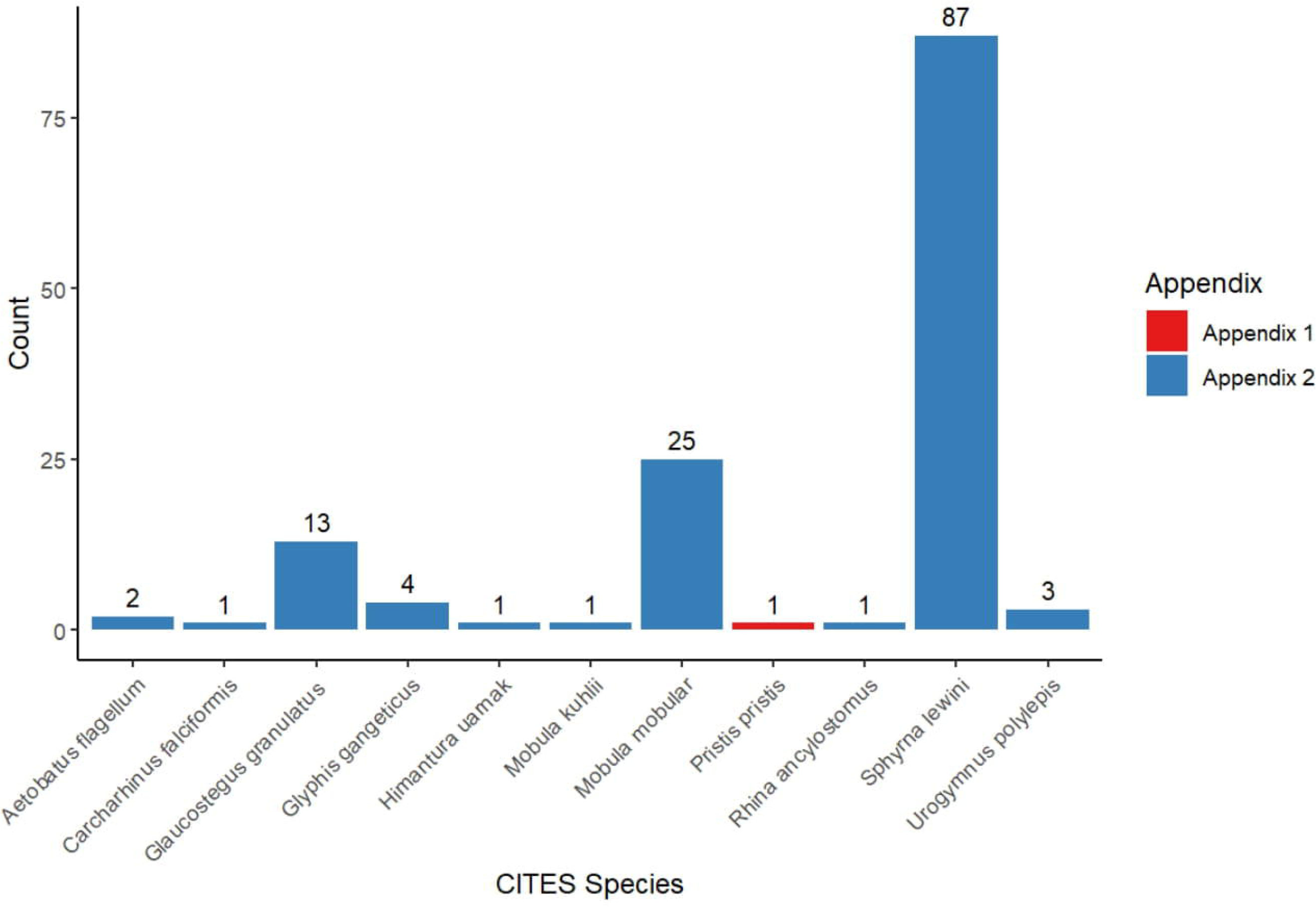
CITES Species and their frequency among the sampled species.

### 3.3. Maturity and sex distribution of sharks, rays, and rhinorays

Of a total of 525 specimens, gender determination of 523 samples was possible. Two specimens were not in good shape to be identified. Among these samples, 65% (64.76%) of the elasmobranchs were female (n=340), 35% (34.85%) of the elasmobranchs were male (n=183), and the rest were unidentified 0.38% (n=2). The total distribution of males vs. females across all species was significantly different (p< 0.0001).

There was a significant difference in the number of sampled shark specimens that were female (72.33%) compared to males (26.69%) (**Figure 8A**). Sex ratio of sampled shark species did differ significantly (χ2 =176.25; P<0.0001). Among the highly sampled species, 78% of *Sphyrna lewini*, 73% of *Carcharhinus leucas*, 71% of *Galeocerdo cuvier,* and 64% of *Carcharhinus sorrah* were female.

**Figure 8:**
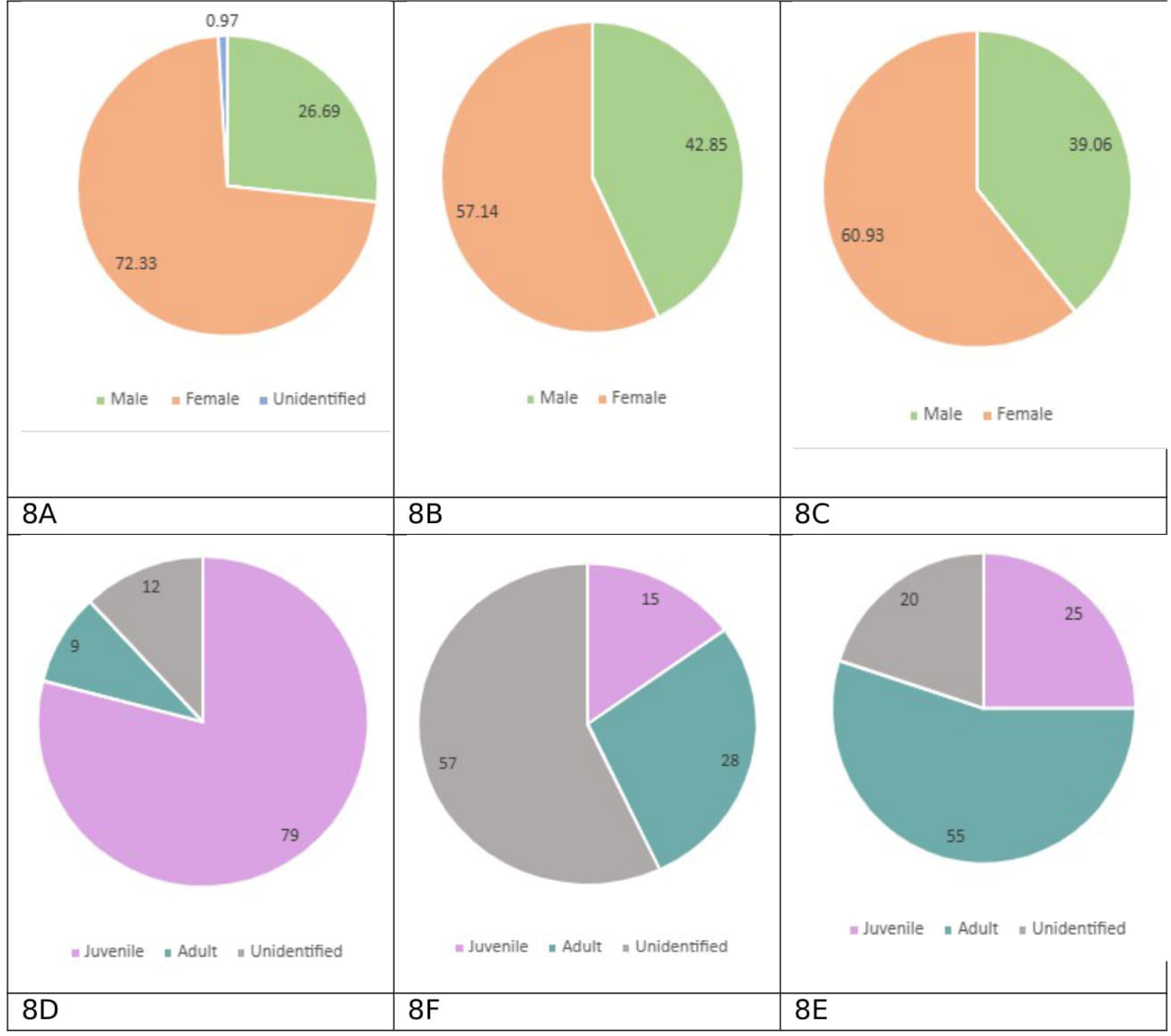
Pie Charts 8A, 8B, 8C showing the gender distribution of Sharks, Rays, Rhinorays and Pie Charts 8D, 8E,8F showing the juvenile vs adult distribution of Sharks, Rays, Rhinorays.

**Figure 9A:**
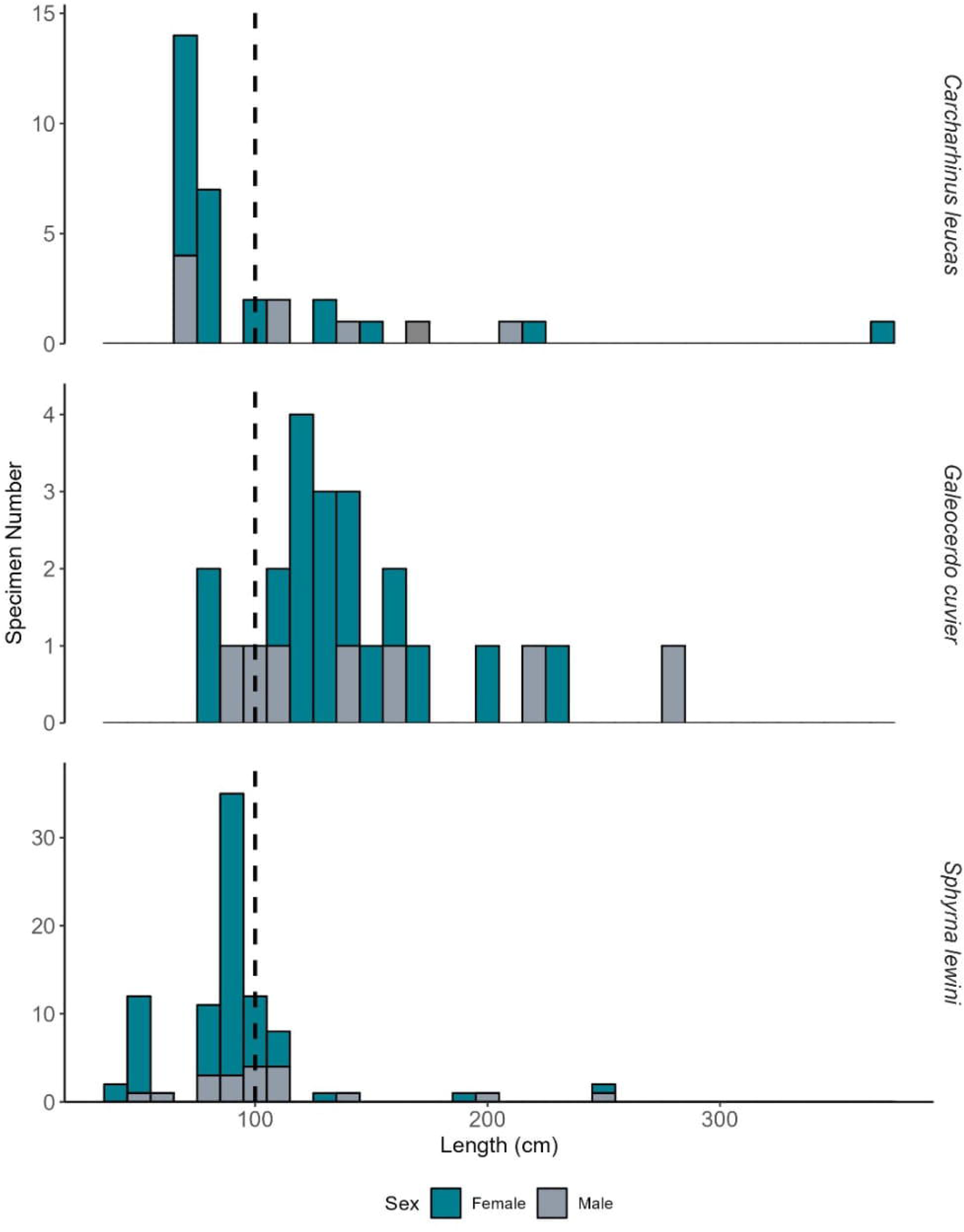
Shows Sharks ***(Carcharhinus leucas, Galeocerdo cuvier, Sphyrna Lewini***) those have large size or have large number of samples along with the male and female ratio.

**Figure 9B:**
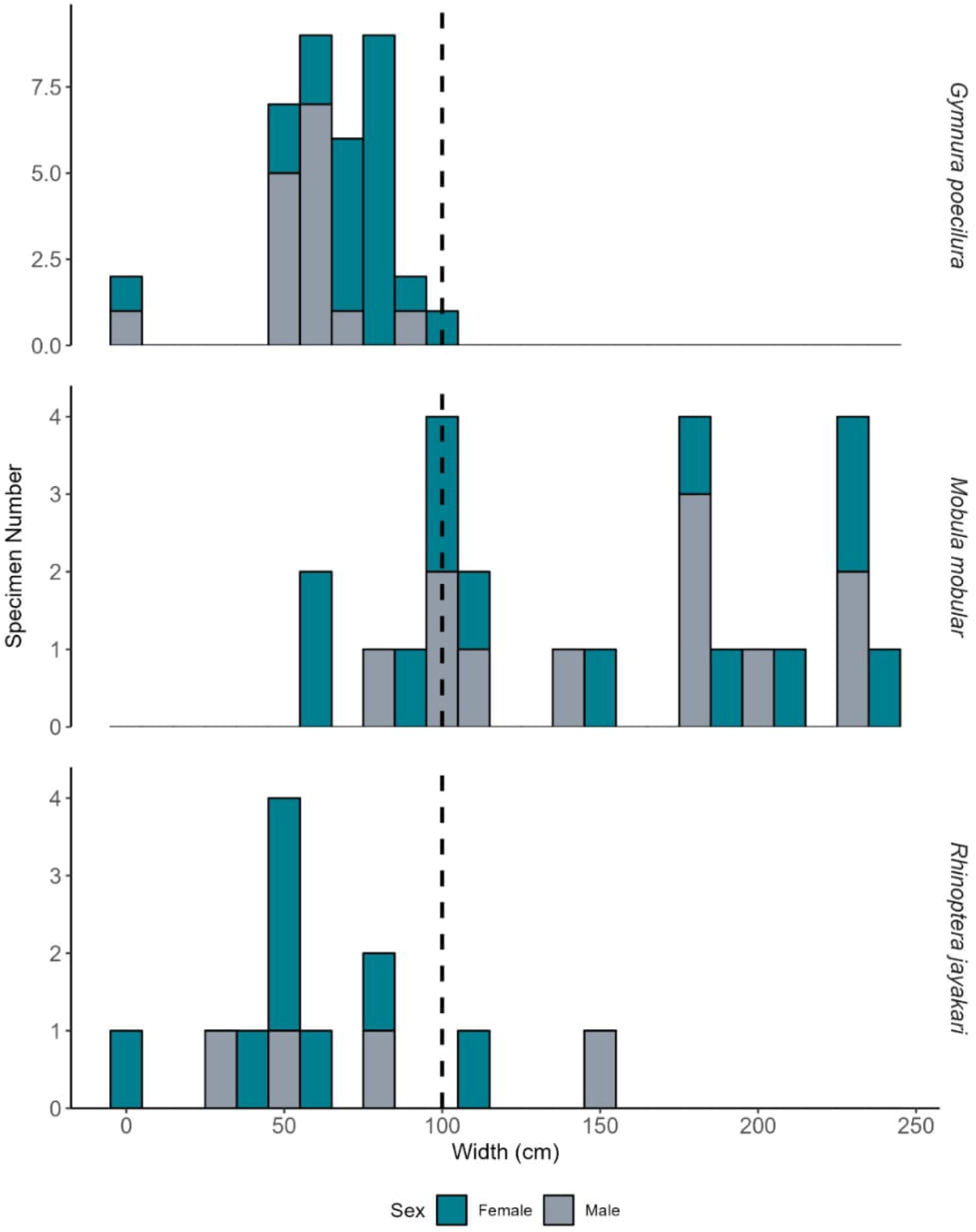
Shows rays ***(Gymnura poecilura Mobula mobular, Rhinoptera jayakari)*** those have large size or have large number of samples along with the male and female ratio.

**Figure 9C:**
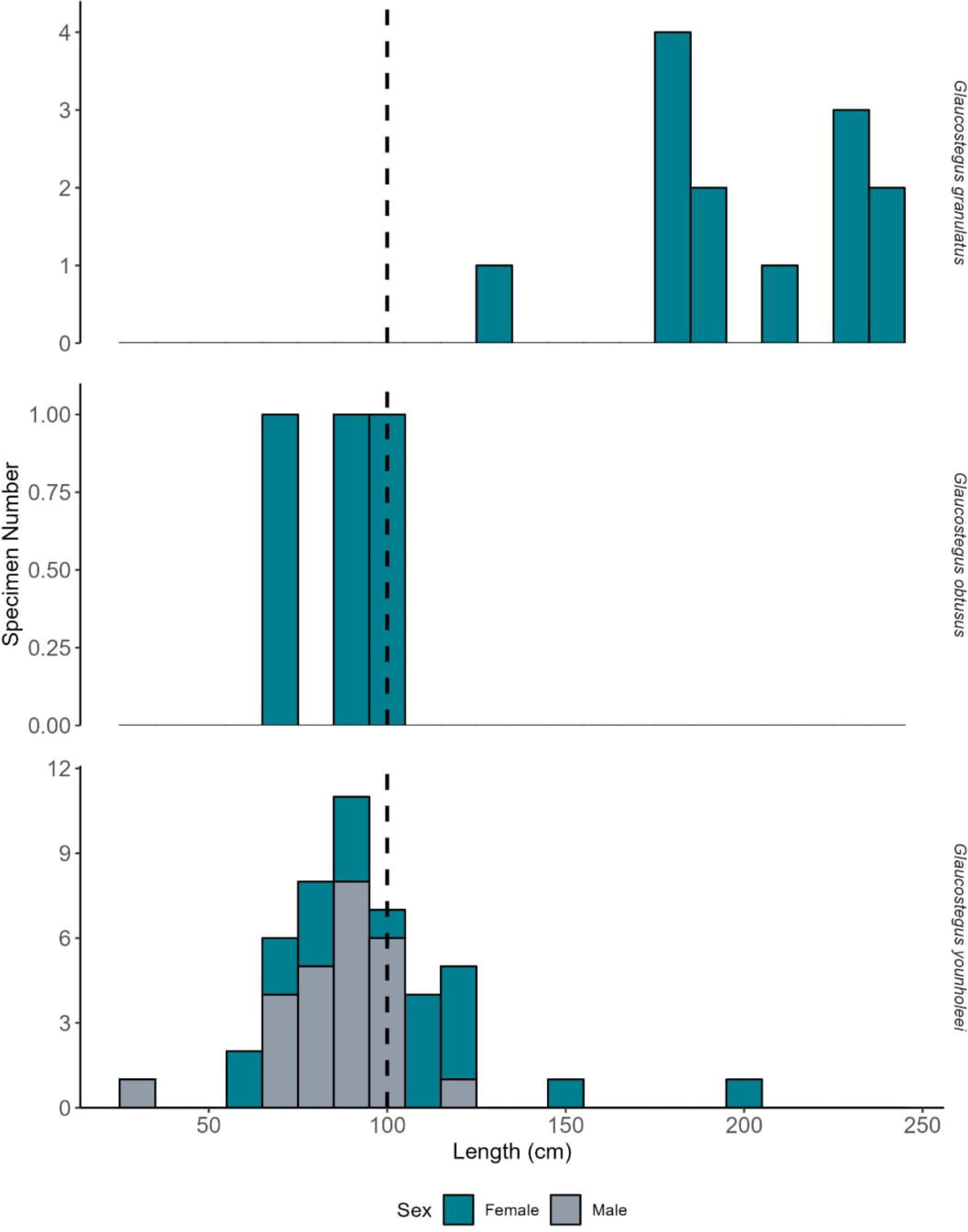
Shows Rhinorays ***(Glaucostegus granulatus, Glaucostegus obtusus, Glaucostegus younholeei***) those have large size or have large number of samples along with the male and female ratio.

The ratio of sampled ray specimens did not differ significantly (χ2 =4.24; P <0.05) (**Figure 8B**).

For multiple species of sharks, rays, and rhino rays, the gender ratio was skewed toward females. For instance, 68 of 87 *Sphyrna lewini*, 17 of the 24 *Galeocerdo cuvier*, 24 of 33 *Carcharhinus leucas*, 8 of 12 *Rhinoptera jayakari* and 13 of 13 *Glaucostegus granulatus* were female (**Table 2**, **Figure 9A and 9B).** Meanwhile, for *Carcharhinus limbatus* the ratio was skewed toward males, with 6 of 11 total male specimens (**Table 2**, **Figure 9A**). For two specimens, specific information on sex was unavailable.

For rays, many samples could not be identified physically or by iNaturalist to the species level, so the sample with genus or family identification was removed from the analysis.

Measurements were analyzed to understand the maturity of the specimens. Among the 525 specimens, length data for sharks and width data for rays were used to determine the maturity or life stage. Juvenile specimens comprised 45% (n=237) of total samples, while adults made up 24% (n=125), and unidentified specimens made up 31% (n=163) of samples.

For shark specimens, 79% (n=185) were juveniles, 9%(n=21) were adults, and 12% (n=27) were unidentified (**Figure 8D)**. Among the mature shark specimens sampled, one of the oldest individuals recorded was for *Galeocerdo cuvier* (Tiger Shark) at 279 cm. The range of the *Galeocerdo cuvier* samples was between 77cm and 279 cm.

For ray specimens, 15% (n=32) were juveniles, 28% (n=59) were adults, and 57% (n=120) were unidentified (**Figure 8E**). Among the rays the most mature individuals were from the following ray species: *Mobula mobular* (Spinetal Devil Ray), followed by *Urogymnus polylepis* (Giant freshwater Stingray). The maximum width found for the *Mobula mobular* was 238.76 cm, and for the *Urogymnus Polylepis,* it was 201cm. This giant freshwater stingray was sampled from the Cox’s Bazar landing station.

For rhinoray specimens, 25% (n=20) were juveniles, 55% (n=44) were adults, and 20% (n=16) were unidentified (**Figure 8F)**. The oldest specimen of a rhinoray recorded was *Glyphis gangeticus* (Ganges River Shark) with a maximum length of 261 cm. The range of this species specimen was between 213 and 261 cm. However, very few samples of Ganges River Shark were sampled during the study.

Similarly, several species had a higher skew towards juvenile specimens, including *Carcharhinus leucas (*Bull Shark), *Sphyrna lewini* (Scalloped Hammerhead), *Carcharhinus brevipinna* (Spinner Shark)*, Galeocerdo cuvier (Tiger Shark)* (**Table 2, Figure 9A),** *Rhinoptera jayakari* (Oman Cownose Ray) (**Table 2**). Conversely, all *Gymnura poecilura* (Longtail Butterfly Ray) (**Table 2**, **Figure 9B**), *Urogymnus polylepis* (Giant Freshwater Stingray) and 12 of 13 *Glaucostegus granulatus* (Granulated Guitarfish) (**Table 2**, **Figure 9C)** were adults. The total number of adults vs. juvenile specimens across all species except *Rhinoptera bonasus* (Cownose Ray) and *Rhinoptera javanica* (Javanese Cownose Ray) was significantly different (p<0.05).

## 4. Discussion

Overfishing, exploitation, and bycatch pose significant threats to elasmobranch populations in the Bay of Bengal, emphasizing the urgent necessity for enhanced management and conservation strategies to avert further extinctions. This study conducted a comprehensive biodiversity assessment of elasmobranchs at two major landing centers in Bangladesh, identifying 29 frequently landed species from a total of 525 specimens collected over two consecutive years during the winter season.

The findings of this study contribute to both global and regional initiatives (Jabado et al., 2017) are aimed at assessing the status of elasmobranchs. Additionally, this research addresses the prevailing gap in scientific literature regarding elasmobranch biodiversity in Bangladesh, as previously highlighted by researchers (Haque et al., 2022). Notably, 86% of the sampled species are classified under IUCN threatened categories, with 38% also listed under CITES, (**Table 2)** thereby reflecting the considerable conservation challenges faced by these species in Bay of Bengal region.

The study reveals that the Bay of Bengal functions as a biodiverse habitat for elasmobranchs; however, fisheries through both targeted and incidental catch are exploiting ecologically critical species at unsustainable rates. Furthermore, this research contributes to mitigating the existing data deficiencies on elasmobranchs (Dulvy et al., 2014; Jabado et al., 2017) by documenting the catch of 29 distinct species. A significant representation of females and juveniles among the landed specimens indicate unsustainable fishing and highlights the critical need for targeted conservation measures (Jaiteh et al., 2017).

### 4.1 Species Distribution and Demography

In our study, elasmobranch distribution in Bangladesh’s two landing sites was dominated by species *Sphyrna lewini* (Scalloped Hammerhead shark) and *Glaucostegus younholeei* (Bangladeshi Guitarfish) being the most frequently recorded. Distinctly, species like the *Pristis pristis* (Largetooth Sawfish) were rare, suggesting localized depletion. The high prevalence of *Sphyrna lewini* in our study samples from both sampling stations raises significant conservation concerns, particularly given its Critically Endangered status on the IUCN Red List. This finding aligns with observations from Indian waters, where *Sphyrna lewini* similarly dominates elasmobranch landings (Thomas et al., 2021), suggesting regional overexploitation patterns.

The capture of a substantial number of juvenile and female elasmobranchs along the coastal waters of Bangladesh suggests the potential existence of nursery grounds in the regions adjacent to Chattogram and Cox’s Bazar. The specimens captured in this study, including *Glaucostegus granulatus* (Granulated Guitarfish), *Sphyrna lewini (*Scalloped Hammerhead) and *Carcharhinus leucas* (Bull Shark) were predominantly females and juveniles. According to CITES, the alarming prevalence of juvenile Scalloped Hammerhead sharks underscores an urgent need for protective measures. Furthermore, the substantial landings of *Glaucostegus younholeei* (Bangladeshi Guitarfish) indicate further investigation into their nursery ground, and the landing of gravid females highlights the necessity for in depth study into their breeding seasons.

### 4.2. Conservation Concerns

The trawling and indiscriminate gear use practices are often employed within the region, and these inadvertently capture numerous non-target species, including a significant number of juvenile elasmobranchs. Our study shows a disproportionate number of juvenile and female indicating fishing in breeding areas and capture of juveniles which are pre-reproductive age and females. This phenomenon hinders the attainment of maturity among these individuals and contributes to the decline of the overall population. The prevalence of juvenile catch signifies the detrimental implications of small mesh sizes and the use of destructive fishing gear, underscoring overfishing’s adverse impacts on population structures.

Our catch data reveal alarming landings of seven Critically Endangered (CE) elasmobranchs: *Sphyrna lewini* 78% females and 96% juveniles, *Glyphis gangeticus*, Guitarfishes, collectively 60% adult females *(Glaucostegus granulatus*, *Glaucostegus younholeei*, and *Glaucostegus obtusus*), *Rhina ancylostomus*, and *Pristis pristis* (**Table 2).** These findings mirror IUCN Red List warnings of >80% population declines regionally (Kyne et al., 2020) with three critical patterns: (1) *Sphyrna lewini* juveniles dominated catches, confirming breeding habitat exploitation; (2) All *Glaucostegus* species showed female bias and gravid females, indicating targeting of reproductive aggregations; and (3) *Pristis pristis* and *Rhina ancylostomus* occurred at reduced frequency compared to previous surveys (Haque et al., 2021), suggesting local extirpation risk. A singular observation of *Pristis pristis* across both sampling seasons reinforces the urgent need to address the pronounced reductions in this population within the Bay of Bengal. Previous literature (Haque et al., 2020) indicates that 25 sawfish were recorded in 2016-17 along the southeastern coast, and this species is recognized to be facing severe declines globally (Dulvy et al., 2021). A comparative analysis of our findings with previous data (Haque et al., 2020) suggests a marked decline in this elasmobranch population within the Bay of Bengal, necessitating immediate conservation actions.

Such trends in landing violate CITES Appendix II protections and Bangladesh’s Wildlife Act 2012 (Revised), demanding immediate gear restrictions (banning gillnets <20 cm mesh), strict enforcement of landing bans for Critically endangered species, and proper implication of management of Swatch-of-No-Ground as a marine protected area. Without intervention, these apex predators’ collapse could destabilize Bay of Bengal food webs within a decade.

Considering the threats posed by global fishing and trade activities, 17 elasmobranch species have been classified under Appendices I and II of the Convention on International Trade in Endangered Species of Wild Fauna and Flora (CITES). Our study documented the catch of 11 CITES-listed elasmobranch species, highlighting significant conservation and management challenges. These species include the *Carcharhinus falciformis* (Silky Shark), *Sphyrna lewini* (Scalloped Hammerhead), *Mobula mobular* (Spinetail Devil Ray), *Mobula kuhlii* (Shortfin Devil Ray), *Aetobatus flagellum* (Longheaded Eagle Ray), *Himantura uarnak* (Honeycomb Stingray), *Urogymnus Polylepis* (Giant Freshwater Stingray), *Rhina ancylostomus* (Bowmouth Guitarfish), *Glaucostegus granulatus* (Granulated Guitarfish), *Pristis pristis* (Largetooth Sawfish), and *Glyphis gangeticus* (Ganges Shark). The persistent capture of these species, despite their CITES Appendix II listings (except *Pristis pristis*, listed under Appendix I), underscores critical gaps in enforcement and species-specific protections in Bangladesh’s fisheries (CITES, 2023).

Under Bangladesh’s updated Bangladesh Wildlife Act, 2012, several critically endangered and ecologically vital elasmobranchs are granted legal protection, prohibiting their capture, trade, or harm. Protected species include the *Glyphis gangeticus* (Ganges shark), *Pristis pristis* (Largetooth sawfish), *Carcharhinus leucas* (Bull shark), *Sphyrna lewini* (Scalloped hammerhead), *Carcharhinus falciformis* (Silky shark), *Carcharhinus amboinensis* (Pigeye shark), and *Galeocerdo cuvier* (Tiger shark). Additionally, the Act safeguards rays such as the *Gymnura spp.* (Butterfly ray), *Mobula spp.* (Devil rays), *Rhinoptera spp.* (Cownose ray), and all species of *Glaucostegus* (Guitarfish) many of which face severe declines due to overfishing and habitat loss. Despite these protections, enforcement remains weak, necessitating urgent measures to prevent further population collapse.

The Bay of Bengal serves as a vital habitat for these threatened species, many of which are highly susceptible to overfishing due to slow growth and low reproductive rates. For instance, *Sphyrna lewini* (Critically Endangered, IUCN) and *Carcharhinus falciformis* (Vulnerable, IUCN) are frequently caught as bycatch in tuna gillnet and longline fisheries, while *Mobula* rays and *Glaucostegus* Guitarfish are targeted for their high-value gill plates and meat in international markets. The presence of juvenile *Glaucostegus gangeticus* (Critically Endangered) and *Pristis pristis* (possibly locally extinct) in landings further indicates the degradation of critical estuarine and mangrove nursery habitats, such as the Sundarbans (Saha et al., 2022).

Despite Bangladesh’s status as a signatory to CITES, inadequate monitoring of elasmobranch trade remains, and the enforcement of protective national laws is lacking. Furthermore, a notable gap exists in awareness among fishers and traders regarding the extant regulations and CITES provisions pertaining to the capture and trade of elasmobranchs, which likely contributes to considerable non-compliance. Urgent measures are needed to align national fisheries policies with CITES obligations, including species-specific catch bans, strengthened trade monitoring, and habitat protection in ecologically sensitive zones. Regional collaboration with neighbouring countries is also essential to curb cross-border illegal trade. Without immediate action, the continued exploitation of these species risks ecological disruptions and the collapse of already declining populations, undermining both biodiversity and the sustainability of small-scale fisheries in Bangladesh.

### 4.3. Challenges

Several individuals belonging to genera such as *Carcharhinus, Glaucostegus, Gymnura, and Rhinoptera* were not identified at the species level, highlighting the challenges associated with on-field morphological identification (Last et al., 2016). This underscores the need for incorporating molecular identification methods alongside morphological techniques to enhance the accuracy of elasmobranch identification and provide a clearer understanding of species diversity within landed catches.

Moreover, the low commercial value of certain elasmobranch species has led to intentionally discarding bycatch, further endangering their populations. Onboard fishing surveys are not widely practiced in the Bay of Bengal, and compliance with logbook entries by fishing crews remains low. The lack of systematic exploratory surveys and limited availability of trained personnel at landing sites further complicated data collection efforts.

Consistent monitoring of this issue is essential for understanding the extent of bycatch discards, as the catch ratios will aid in elucidating the current population structure of various species. Comprehensive data on catch per unit effort, the range of trawlers, and multi-seasonal assessments are vital for identifying species-specific nursery grounds. Collecting year-round data, alongside seasonal and biotic-abiotic water data, will yield deeper insights into habitat utilization and potential nursery ground dynamics.

### 4.4. Further Directions

Overfishing and exploitation pose significant challenges to elasmobranch conservation efforts. These initiatives often fail when fishing bans or restrictions are implemented without offering economic alternatives to affected communities. In Bangladesh and the larger Bay of Bengal region, where small-scale fishers rely heavily on shark and ray fisheries, a punitive approach such as strict bans without compensation results in non-compliance, resistance, and even increased illegal fishing. So economic alternative, economic incentive drive compliance and gender inclusivity will be more sustainable.

To enhance elasmobranch conservation in Bangladesh, future initiatives should focus on live release programs that train fishers in proper handling techniques to reduce post-release mortality. Coupling these programs with incentives such as direct payments or community recognition may further encourage participation.

Awareness workshops for fishing communities should emphasize the ecological significance of sharks and rays and the benefits of sustainable practices. Additionally, introducing alternative fishing gear, like circle hooks and bycatch reduction devices, can help mitigate accidental catches while maintaining fishery productivity.

A collaborative approach that merges fishers’ knowledge with scientific research is vital for developing practical and economically viable conservation measures. By integrating science-based interventions, socioeconomic incentives, and effective policy enforcement, Bangladesh can decrease elasmobranch mortality, protect marine biodiversity, and support coastal livelihoods, creating a replicable model for other tropical small-scale fisheries.

## 5. Conclusions

In conclusion, this study highlights the urgent need to address the knowledge gap in elasmobranch landings in Bangladesh, where data scarcity hinders effective conservation. The findings emphasize the importance of strengthening monitoring efforts, particularly for protected and threatened species, while advocating for species-specific research to identify critical habitats like breeding and nursery grounds in the Bay of Bengal. Sustainable management requires a multi-stakeholder approach, engaging fishers, policymakers, researchers, and local communities to implement science-based conservation strategies. Additionally, strict enforcement of existing laws, such as the Bangladesh Wildlife Act 2012, alongside targeted fishing bans and habitat protection, is crucial to prevent further population declines. Proactive measures must be taken now to ensure the long-term survival of these ecologically and economically important species.

## Authorship Contribution Statement

Fahmida Khalique Nitu helped with the study design, field supervision, organized and analysed the data, and wrote the first draft and revised manuscript version. Dilshad Tasnima collected and organized the data and uploaded it to iNaturalist. Sultan Ahmed collected the data and helped in the formation of the map. Kutub Uddin helped with the project design, project supervision and manuscript review. Shaili Johri designed and supervised the study and contributed to writing and editing of the manuscript.

## Appendix A. Supplementary files

Table 3

## Data availability

Data will be made available on request.

---

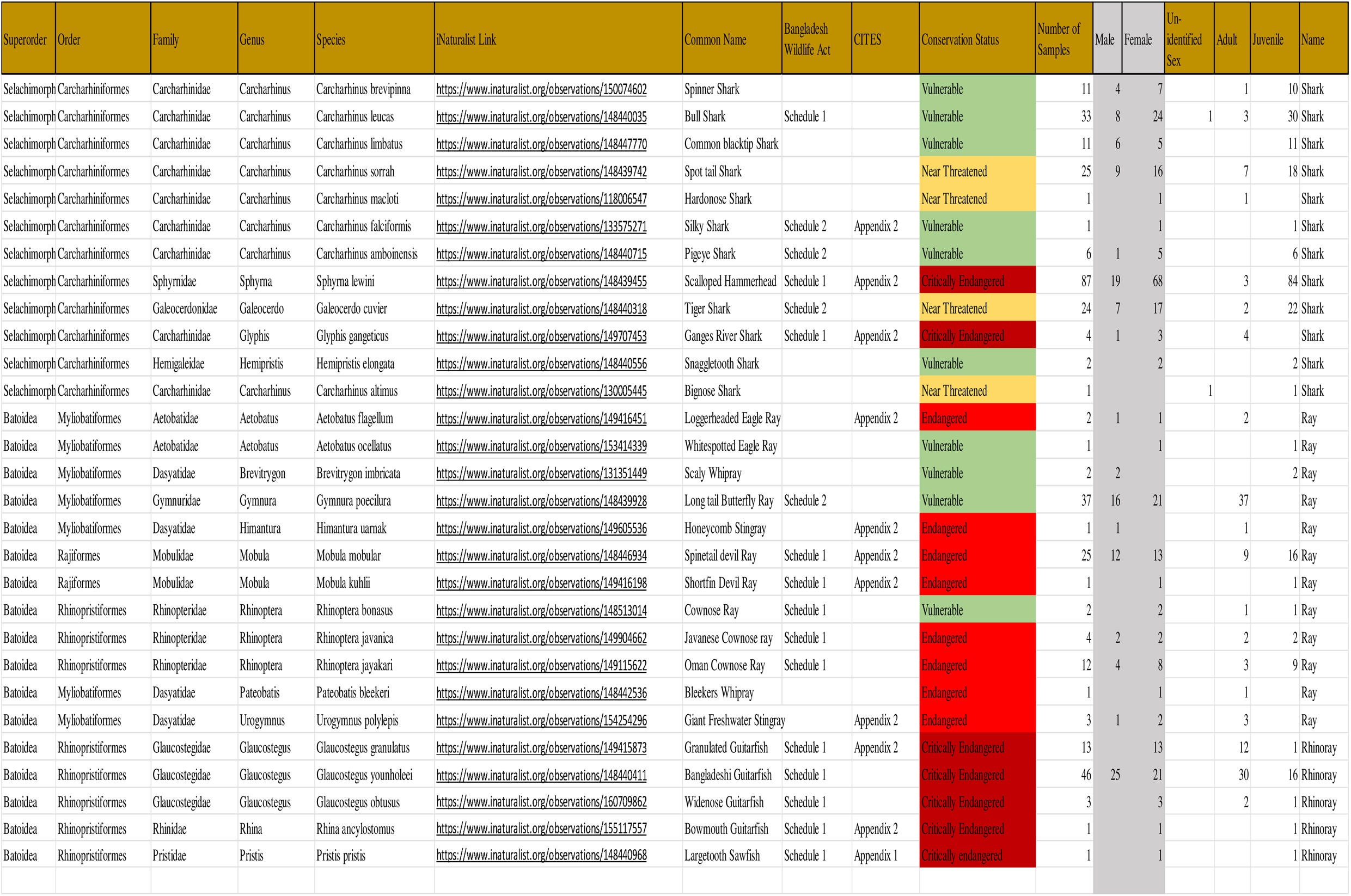

